# Trial-History and Spatial Biases Shape Perceptual Decision-Making in Archerfish

**DOI:** 10.64898/2026.06.24.734265

**Authors:** Ori Hendler, Kasturi Mondal, Nimrod Dushnik Shamir, Svetlana Volotsky, Maoz Shamir, Ronen Segev

**Author notes:** Equal contribution (Ori Hendler, Kasturi Mondal).

## Abstract

Identifying conserved principles of animal decision-making requires comparisons across evolutionarily distant species with divergent neuroanatomy. Yet teleost fish, an evolutionarily distant vertebrate lineage lacking a laminated cortex, are rarely studied using behavioral paradigms directly comparable to those used in mammals and birds. Here, we bridge this gap by exploiting the unique hunting behavior of archerfish to investigate determinants of decision-making. We trained archerfish to report their choices by shooting a jet of water at the selected visual target in a series of two-alternative choice tasks. We found that archerfish decisions are influenced by trial-history and spatial-biases, remarkably similar to those observed in mammalian and avian models. Specifically, subjects exhibited an outcome-history in the form of stay/switch behaviors, alongside a stimulus-history bias, in which they repeatedly selected the same shape across trials. Yet, unlike rodents, archerfish maintained stable behavior during unrewarded probe trials. Thus, our findings reveal that several fundamental determinants of choice previously characterized primarily in mammals are also present in teleost fish, suggesting that important components of vertebrate decision-making are conserved despite substantial divergence in brain organization and ecology. Simultaneously, the distinct response of archerfish to reward omission highlights species-specific differences in how animals adapt to changing environmental contingencies.

## INTRODUCTION

Over the past decade, studies of animal decision-making have proven fruitful in illuminating several variables that can shape choice [1–4]. In particular, controlled behavioral studies across species, including humans [5], non-human primates [6], and rodents [7,8], have been instrumental in identifying shared determinants of decision-making [9,10]. One such determinant is termed spatial bias, in which choices are skewed toward a specific location. For example, in studies of non-human primates, subjects exhibit idiosyncratic preferences for spatial locations, despite all locations being equally rewarded [11–14]. Likewise, rodents performing discrimination tasks exhibit spatial biases, which may become more pronounced with task difficulty [15–17]. Another major determinant is trial-history, whereby choices can depend not only on the current stimulus but also on recent stimuli and previous outcomes [18,19]. Such biases have been documented across species, including humans [20,21] and rodents [22,23], and have been shown to persist even in well-trained subjects [24,25].

Identifying conserved principles of decision-making benefits from comparisons across species that diverged early in evolutionary history and face distinct ecological demands [26,27]. Teleost fishes, lacking a mammalian-like cerebral cortex, provide a valuable comparative model for identifying principles of cognition and decision-making that may be shared across divergent vertebrate lineages [28,29]. However, decision-making in teleosts has rarely been studied using experimental paradigms directly comparable to those employed in mammals. This methodological gap makes it difficult to determine whether well-established determinants of mammalian choice similarly influence decision-making in teleosts, and, if so, to what extent.

Archerfish provide a unique opportunity to address this question [30]. In nature, archerfish hunt by shooting jets of water at prey above the water surface. In the laboratory, they can be trained to shoot at visual stimuli presented on a screen [31], providing a discrete behavioral report of their choice on each trial. This allows archerfish decisions to be quantified using paradigms directly comparable to those used in mammals, such as reaching in primates or lever pressing in rodents, facilitating cross-species comparisons of decision-making. Over the past two decades, archerfish have been used to investigate a broad range of visual and cognitive abilities, including orientation saliency and pop-out search [32], inhibition of return [33], object recognition [34], and motor adaptation [35]. However, their decision-making determinants have not been addressed.

In the present study, we investigated decision-making in archerfish. We trained subjects on a series of two-alternative non-forced choice tasks and modeled their trial-by-trial choices using dynamic generalized linear models [2]. This approach allowed us to track how distinct choice determinants shift during reinforcement learning. We show that target selection exhibits slowly drifting spatial biases, and that choices are influenced by stimulus-history and outcome-history, with transient stay/switch behaviors emerging during the experiment. Ultimately, teleost fish share these choice determinants with diverged vertebrate lineages, suggesting that key aspects of decision-making have been evolutionarily conserved, transcending divergences in brain structure and habitat.

## RESULTS

To investigate decision-making, we trained archerfish in a two-alternative non-forced choice task in which two shapes, a target and a non-target, were presented simultaneously on a monitor (Experiment 1, Fig. 1a, see Methods). Selection of the target was rewarded with a food pellet, whereas selection of the non-target was not rewarded. On rare probe trials, reward was omitted regardless of the fish’s selection (Fig. 1a). On the shape task, the stimuli consisted of spider and ant images (see Methods).

**Figure 1.**
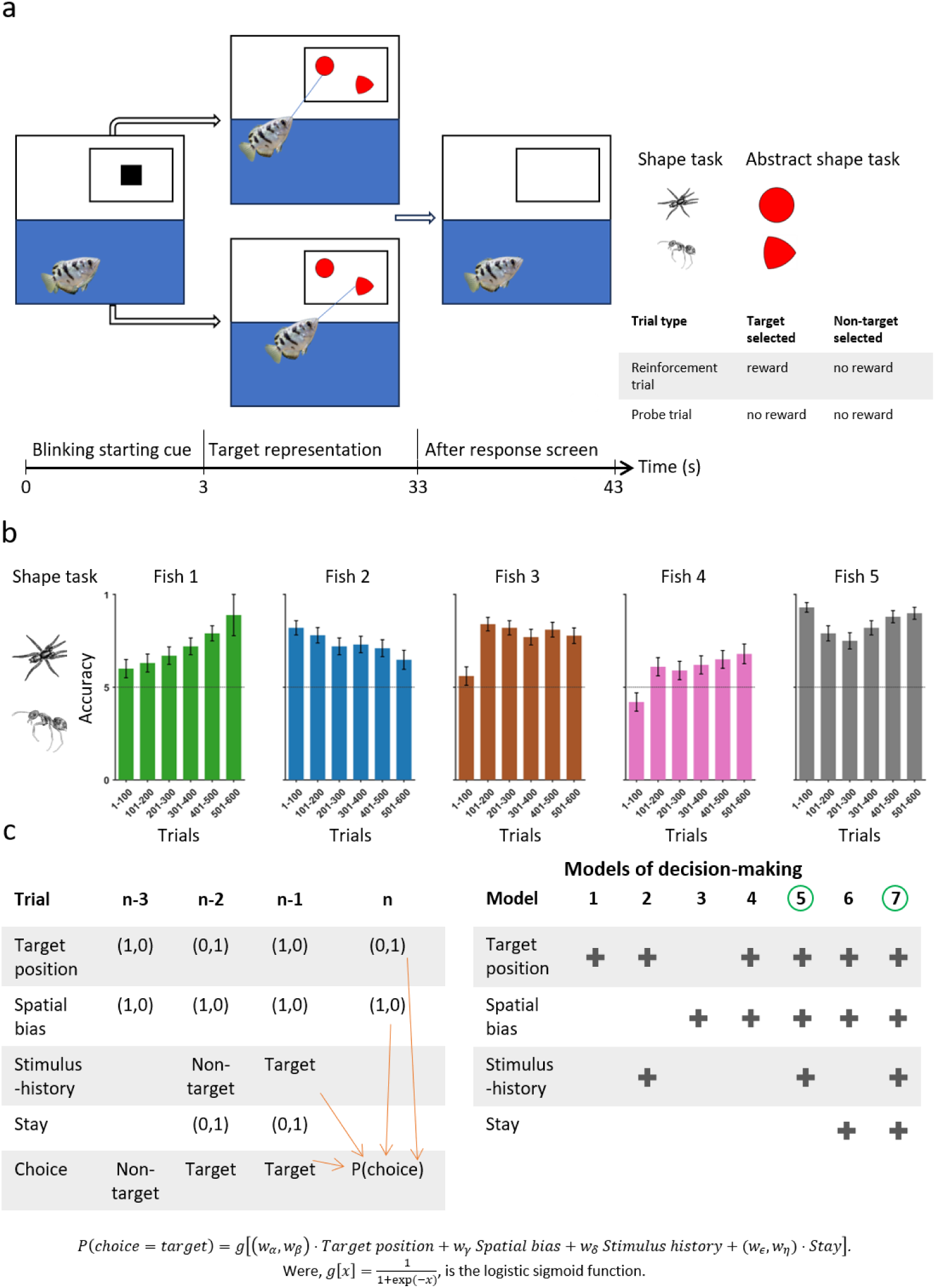
Task illustration, behavioral accuracy and modeling framework. **(a)** Trial structure. Each trial began with a 3 s blinking start cue, followed by a 30 s stimulus presentation phase in which a target and a non-target were displayed simultaneously on the screen above the tank. The fish indicated its choice by spitting a water jet at one of the two stimuli. The trial ended with a 10 s blank screen, giving a total trial duration of 43 s. The schematic at the top right shows the two task variants. In the shape task, the stimuli were images of spiders and ants. In the abstract shape task, the stimuli were two red abstract shapes of equal area. The table at the bottom right shows the two trial types. In reinforcement trials, choosing the target was rewarded and choosing the non-target was not. In probe trials, reward was omitted irrespective of choice. **(b)** Presents the behavioral accuracy of Fish 1-5 during the two-position shape task, Experiment 1. Bars show mean accuracy ± SEM in consecutive bins of 100 trials, and the dashed line marks chance level. **(c)** The table on the left illustrates the decision-making variables over an example sequence of trials, from trial n−3 to trial n. orange arrows illustrate the influence of these variables on the probability of selecting the target, P(choice). The table on the right lists the seven models tested on the task, M1 to M7, which differ in the subset of decision-making variables they include. A plus sign marks a variable that is included in a model. Green circles indicate best fitting models. The equation below the tables is the logistic generalized linear model that links the decision-making variables to choice probability, where g[·] is the logistic function.

### Archerfish display complex dynamics in behavioral accuracy during a shape discrimination task

We found that rather than a simple monotonic increase in success rate until a plateau was reached, behavioral accuracy across the five fish tested on the shape task was heterogeneous and exhibited complex dynamics (Fig. 1b). Some fish showed a gradual rise in accuracy across trials (Fish 1 and Fish 4), while others displayed the opposite pattern, with accuracy declining progressively over the course of the experiment (Fish 2). Some fish followed a non-monotonic trajectory, with accuracy decreasing (Fish 3) or increasing (Fish 5) later in the experiment. Such complex trajectory profiles raise the question of what decision-making determinants drive choice on this task.

### Modeling decision-making with dynamic generalized linear models

To identify the factors influencing choice behaviour, we modelled target versus non-target selection using a dynamic generalized linear framework (see Methods). Target position was included as a predictor of choice, with separate terms for each possible target position. To capture fish-specific spatial preferences, we included a spatial bias term. We also incorporated two history-dependent variables to capture the influence of previous trials on current choice: a stimulus-history term, capturing the tendency to repeatedly select a particular shape, and an outcome-history term, capturing the tendency to either return to a previously rewarded location (win-stay) or remain at a previously unrewarded location (lose-stay). These variables were combined in an inference model that estimated, on each trial, the probability that the fish would select the target shape according to:

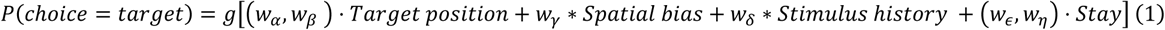

Where, *g*[⋅] is the logistic function. Figure 1c (left panel) illustrates an example sequence of trials (n−3 to n) and the influence of the different choice determinants, indicated by orange arrows, on the probability of choice on trial n.

### Best fitting model across all fish includes both spatial and temporal determinants of choice

We compared seven models (M1–M7, see Methods) that differed in the subsets of decision-making variables included in equation (1) (Fig. 1c, right panel). Across fish, the model best describing the data (based on the Bayesian Information Criterion, BIC, see Methods) was either M5 or M7, indicated by green circles, both of which included spatial and temporal determinants of choice. Per-fish comparisons of all models for the 14 fish are provided in Supplementary Figs. S1–S5.

### Target position, not spatial bias, dominates choice behavior

The behavioral accuracy of three representative fish (black) and their corresponding model fits (cyan) are illustrated in Fig. 2a–c (see Supplementary Fig. S1 for model comparisons). Accuracy when the target was in positions one and two is shown in dark gray and light gray, respectively. Notably, performance was highly dependent on the target’s position. Based on the Bayesian Information Criterion, we found that model M7, which includes all variables, was the best fitting model for all 5 fish in this task (see the BIC comparison in Supplementary Fig. S1).

**Figure 2.**
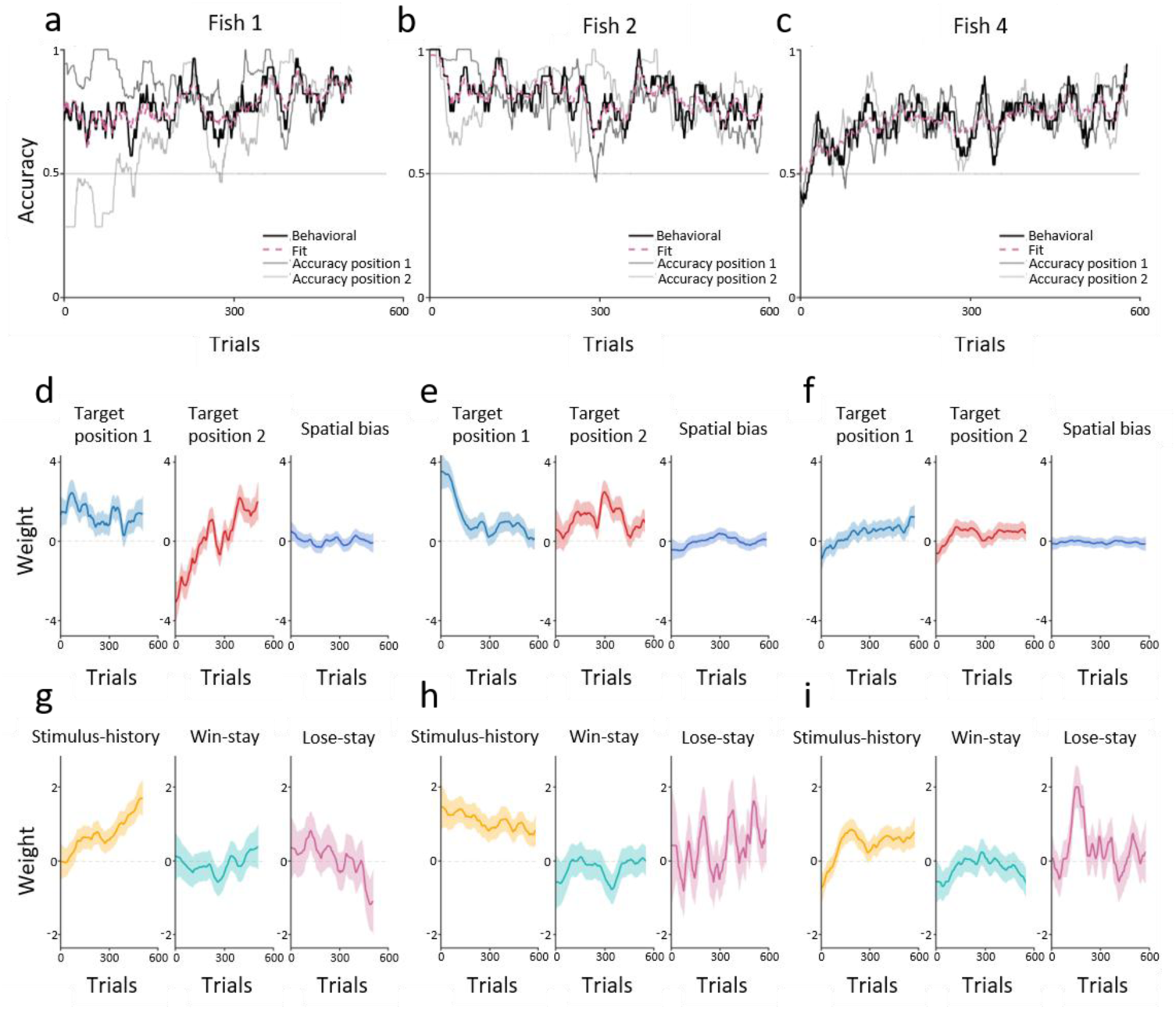
Accuracy and decision-making variables in the two-position task. **(a–c)** Behavioral accuracy over trials is shown for three representative fish: Fish 1, Fish 2, and Fish 4, respectively. Behavioral accuracy is shown in black, and the best-fitting model is shown as a cyan dashed line. Accuracy for the target in position 1 and position 2 are shown in dark gray and light gray respectively. The dashed gray line marks the chance level. The number of trials for position 1 and position 2 are not necessarily equal, because this was a non-forced choice task. In addition, not all fish completed the same number of trials, as detailed in the Methods. (**d**) The three spatial decision-making variables for Fish 1; namely, target position 1, target position 2, and spatial bias. Solid lines show the posterior MAP estimates, and shaded bands show the ±1.96 posterior SD. (**e**, **f**) The same as panel d, for Fish 2 and Fish 4, respectively. (**g**–**i**) The three history decision-making variables for the fish: stimulus-history, win-stay, and lose-stay. Conventions are as in panels d to f.

We examined two spatial decision-making variables: target position and spatial bias. Throughout the experiment, the weights for the two target positions followed markedly different trajectories (Fig. 2d–f). In contrast, the spatial bias weight remained relatively fixed in time with confidence intervals overlapping zero for all three fish (Fig. 2d–f, rightmost panels). Hence, choice was primarily driven by target position and not by a simple spatial bias.

### Decision-making is influenced by trial-history

Beyond the spatial variables, the best-fitting models for all fish also incorporated trial-history variables (Fig. 2g– i). Specifically, we defined a stimulus-history variable (see Methods) to quantify the subjects’ tendency to reselect the same visual shape they had chosen in the previous trial. For all fish, the weight of this stimulus-history variable remained positive throughout most of the experiment (Fig. 2g–i, leftmost panels). We confirmed that stimulus-history reflected a behavioral tendency beyond what accuracy alone would produce by comparing the binary choices to shuffled data. In 7.3%, 10.2% and 8.3% of the experiment, Fish 1, 2 and 4, respectively, had significantly (one-sided binomial test, *p* < 0.05) more repetitions of a shape, than for the shuffled data. Critically, this means that many of the errors during the experiment were driven by repetition of the previous selection rather than by failure to perceptually discriminate the target from the non-target shape.

The effects of the outcome-history variables, win-stay and lose-stay, were heterogeneous across fish. The win-stay weight, which captured the tendency to return to the same position following a rewarded trial, remained close to zero with confidence intervals overlapping zero in Fish 1 and Fish 2 (Fig. 2g–h, middle panels), but in Fish 4 it drifted to clearly negative values during the last portion of the experiment (Fig. 2i, middle panel), indicating that this fish progressively avoided positions where it had just been rewarded. In Fish 1, a negative lose-stay weight developed by the end of the experiment (Fig. 2g, rightmost panel), indicating a growing tendency to switch positions after errors. By contrast, Fish 2 exhibited high lose-stay around trials 200-300 and again around trials 400-500 (Fig. 2h, rightmost panel). Hence, we find idiosyncratic outcome-history in archerfish.

The same analysis was applied to all five fish on this task (see Supplementary Fig. S2). Across all fish, target position and trial-history had a non-stationary impact on choice.

### The effects of position and trial-history on choice, generalize to a four-position task and to abstract stimuli

Given the prominent role of target position on choice, we ran a follow-up experiment in which the shape task was extended from two to four possible target positions: the target and non-target could now appear anywhere on the screen, with the constraint that they could not be in the same quadrant (Experiment 2, see Methods). Spiders and ants represent ecologically relevant prey for archerfish. To test whether this influenced the decision variables observed previously, we conducted a second version of the four-position task using abstract shapes, a circle and a morphed triangle (Experiment 3, see Methods). Figure 3a,b show the behavioral accuracy for the shape task and the abstract shape task, respectively.

**Figure 3.**
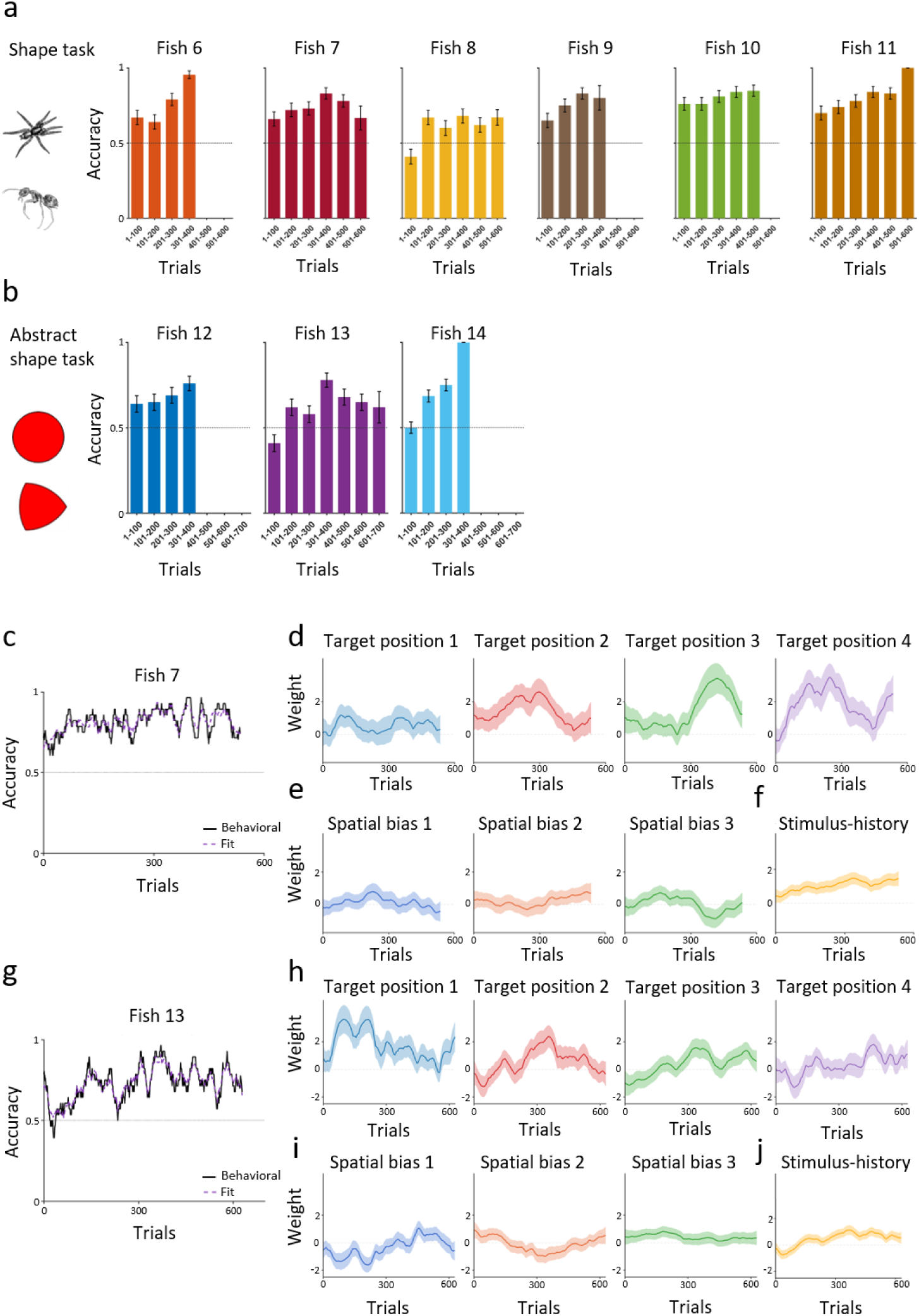
Accuracy and decision-making variables in the four-position task. **(a)** Behavioral accuracy for the six fish that performed the four-position task, Fish 6 to 11. Data are shown as mean accuracy ± SEM in consecutive 100-trial bins. The dashed line marks the chance level accuracy. Not all fish completed the same number of trials, as detailed in the Methods. **(b)** Behavioral accuracy across trials for Fish 12-14, which performed the abstract shape task in Experiment 3. Conventions as in panel a. **(c–f)** Model fit and decision-making variables for Fish 7. **(c)** The black line shows the accuracy of the fish across trials, and the dashed purple line shows the fit of the best model, M5. This model includes three types of decision-making variables: target position, spatial bias and stimulus-history. Panels d-f show the weight of each variable across trials, that is, how strongly it influenced choice. **(d)** Weights of the four target position variables. A higher weight means the fish was more likely to choose the target when it appeared at that position. **(e)** Weights of the three spatial bias variables. Positive weights indicate a preference for a position and negative weights an avoidance, regardless of where the target was. **(f)** Weight of the stimulus-history variable. A positive weight means the fish tended to choose the same shape as on the previous trial. In panels d-f, solid lines show the posterior MAP estimates and shaded bands show ±1.96 posterior SD. **(g–j)** Same as c to f, for Fish 13.

The accuracy curves in Experiment 2 and Experiment 3 varied considerably across individuals. Some fish demonstrated a gradual increase in accuracy over the course of training (Fish 6, 10, 11, 12 and 14), while others maintained a relatively flat trajectory (Fish 8), or exhibited an initial increase followed by a subsequent decline (Fish 7 and 13). These findings demonstrate that coarse, aggregated measures of accuracy are insufficient to capture individual performance and that interpreting these results requires an examination of the trial-by-trial choices.

We extended the dynamic generalized linear model framework to the four-position task as detailed in Methods (see Supplementary Figs. S3–S5). Here we highlight two representative patterns of behavior: Fish 7’s behavior on the shape task (Fig. 3c) and Fish 13’s behavior on the abstract shape task (Fig. 3g). In both fish, the four target position weights, one for each of the four possible target locations, had a strong and non-stationary effect on choice (Fig. 3d,h). Spatial biases also impacted decision-making, shifting over long timescales of ∼200 trials (Fig. 3e,i). In terms of the history-dependent variables, the stimulus-history remained positive, even when the overall behavioral accuracy declined (Fig. 3f,j). All subjects had significantly (one-sided binomial test, *p* < 0.05) more repetitions of a shape, than for shuffled data (see Methods). We also found that the best fitting model across all fish was model M5, which included both spatial and trial-history variables (see the model comparison in Supplementary Fig. S5). Together, these results demonstrate that the effects of target position and trial-history on choice, were not specific to the two-position task or to ecologically relevant stimuli but were general features of decision-making in archerfish.

### Trial-history variables improve model fit

To quantify the general contribution of trial-history to choice, we compared models with and without history terms for each of the 14 subjects individually. As shown in Fig. 4a, including history variables consistently improved the goodness of fit and yielded lower BIC values (Wilcoxon signed-rank test, p < 0.05). We also note that, while trial-history is found to be a major determinant of choice across all subjects, the directionality of the stay/switch contingencies and the magnitude of stimulus-history are idiosyncratic.

**Figure 4.**
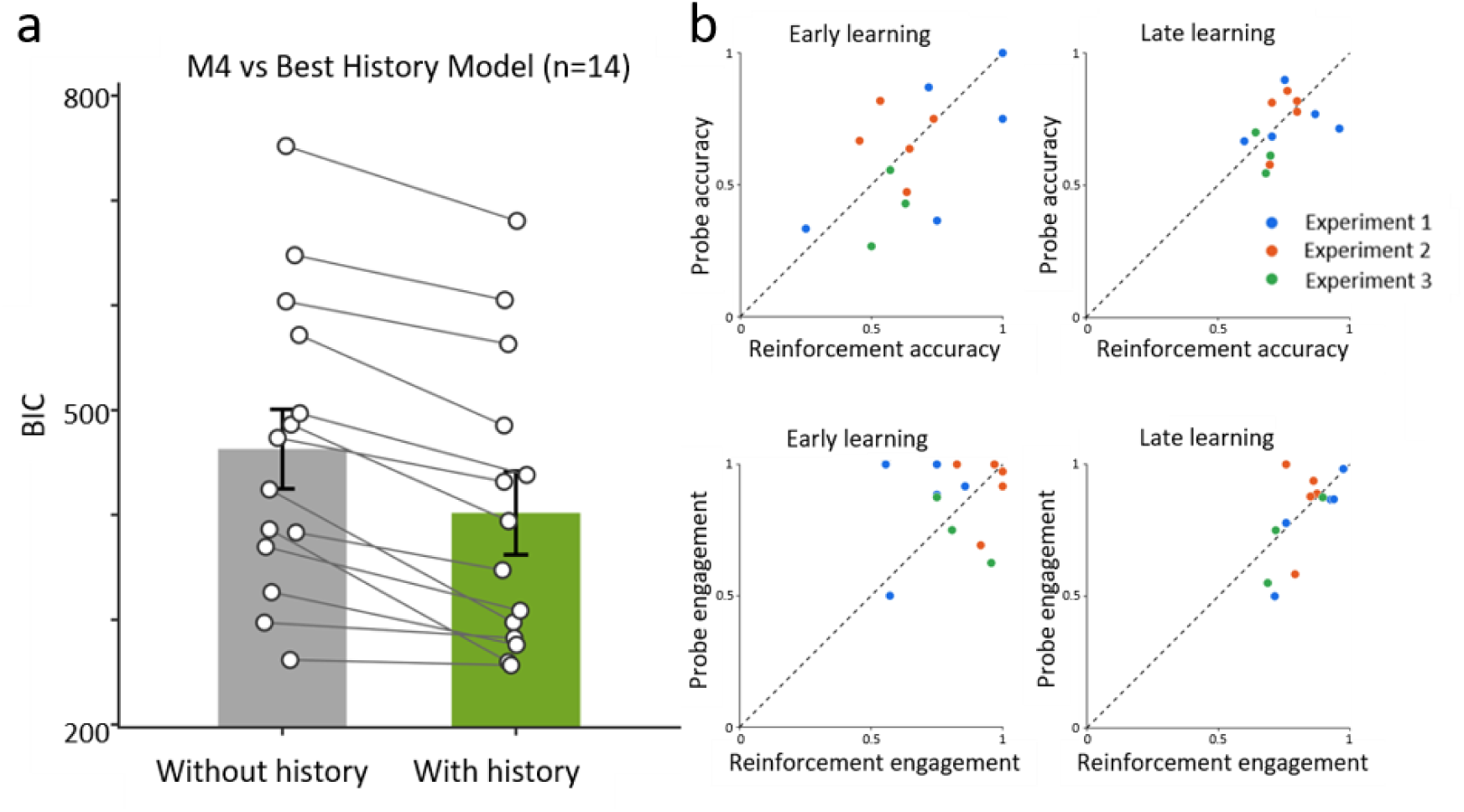
Trial-history improves model fit, and performance in probe and reinforcement trials was similar. **(a)** Model comparison by BIC, where lower values are a better fit. For all 14 fish, the model including trial-history variables had a lower BIC than the model without them. For each fish, model M4, which included only target position and spatial bias, was compared with the best-fitting history model: the lowest-BIC model that also included at least one of stimulus-history, win-stay or lose-stay. Open circles show individual fish, thin lines connect the two values for each fish, and bars show the mean ± SD across fish. **(b)** Probe trials vs. reinforcement trials, split into the early learning phase, left column, and the late learning phase, right column. Top row, accuracy on the probe trials plotted against accuracy on the reinforcement trials. Bottom row, engagement on probe vs. reinforcement trials. Each dot represents the data for one fish and dashes show the identity line.

### Reward omission does not alter choice behavior

Next, we tested whether removing the reward during the probe trials affected choice behavior. For this purpose, we compared accuracy and task engagement on reinforced versus probe trials separately for the early and late phases of learning (Fig. 4b); one fish was excluded (see Methods). We tested equivalence to determine whether probe and reinforced performance fell within a predefined margin of each other. Across all fish, probe and reinforced performance were statistically equivalent in both accuracy (early TOST p = 0.033, late p < 0.01) and engagement (early p = 0.029, late p = 0.001). Hence, archerfish choice behavior was robust to brief omissions of reward.

## DISCUSSION

Across three experiments, we characterized core elements of archerfish decision-making, demonstrating that trial-history shapes choices in two distinct ways. First, choices were influenced by outcome-history if the form of a transient stay/switch behavior. Second, choice was shaped by a stimulus-history bias, whereby fish tended to repeat their selection of a specific shape from one trial to the next. In the spatial domain, choices were governed by the conjunction of target shape and position rather than by shape alone. Notably, this influence was non-stationary, drifting on a slow timescale of approximately 100 trials. Furthermore, a simple spatial bias, a baseline preference for a specific location irrespective of shape, emerged in Experiments 2 and 3, but was absent in Experiment 1.

The determinants of choice we identified in archerfish closely resemble those reported in other vertebrate lineages that diverged later in evolutionary history. Such resemblance strengthens the notion that multiple fundamental choice determinants are shared across vertebrate taxa. In humans and non-human primates, choice behavior is influenced by spatial biases and by the history of previous choices and outcomes [36–38]. For instance, a preference for the conjunction of shape and position, predominantly when a task is easy, has recently been reported [39]. In addition, a general spatial preference has also been shown in humans and non-human primates [12,40]. In rodents, a major part of experimental design and training is concerned with the impact of spatial and trial-history biases and the means to mitigate them [41,42]. In particular, stimulus-history has been shown to modulate choice in rodents [43] in a manner akin to the findings reported in our study. Finally, much like the archerfish in our study, well-trained avian species have demonstrated persistent trial-history biases that consistently influenced their choices [44,45].

Despite these similarities, the temporal dynamics of these determinants differed across species. In rodents, the impact of both task-irrelevant trial history and early spatial biases typically declines with continuous training [2,8,24]. In contrast, archerfish exhibited episodic trial-history effects and a complete absence of spatial bias in Experiment 1. When spatial biases did emerge in this study, during the four-position tasks, Experiments 2 and 3, these biases dynamically fluctuated across trials rather than systematically diminishing over time.

It is reasonable to assume that the determinants of choice identified here are widespread among teleost fishes [46]. If so, behavioral accuracy in a given task should not be interpreted as a direct proxy for psychophysical capacity. Experimental studies in fish may therefore, where possible, account for and mitigate trial-history and spatial biases.

In contrast with rodent studies, where shifts in reward contingencies typically trigger marked, immediate adjustments in subsequent behavior [24,47], we found that brief reward omission did not measurably alter either choice accuracy or task engagement in archerfish (Fig. 4b). One possible interpretation is that archerfish may exhibit lower behavioral flexibility. This possibility is consistent with previous work, which has established a role for the mammalian cortex in supporting flexible switching between behaviors [48,49]. Alternatively, the absence of a behavioral response to reward omission may indicate that archerfish (unlike rodents, see for example [50,51]) are a somewhat non-explorative species, thereby maintaining stable choice behaviour despite short-term changes in external contingencies. These possibilities make different predictions about how fish respond when a previously learned contingency is reversed. Reversal-learning paradigms, such as those used in rodents [52], could therefore provide a test of whether the observed stability reflects low behavioral flexibility or stable behaviour in the face of changing reward conditions [53,54].

The neural substrates of these choice determinants in teleost fish remain unknown, because the circuit-level machinery of reward-based choice has been far less characterized in teleost fish than in mammals, where trial-history signals have been recorded in posterior parietal and prefrontal cortex [7,8,37,55]. Teleosts possess candidate circuitry, an ascending dopaminergic projection from the posterior tuberculum to the subpallium [46,56], proposed as the analog of the mammalian mesostriatal pathway (although this homology remains debated [56,57]) which serves as the primary neural substrate for value-based, and motor-executable decision-making. Another area of interest is the habenula-raphe pathway that encodes aversive expectation and has been shown to support outcome-dependent behavioral updating [58]. Ultimately, we hope the current study sets the stage for future physiological work to probe the single-trial correlates of trial-history biases and reward prediction error signals in teleost fish.

In the current study, we report that trial-history shapes perceptual decision-making in archerfish. Interestingly, a recent study of larval zebrafish automatic motor behavior [59], showed that previous sensory stimuli bias subsequent motor responses away from the preceding stimulus, whereas previous responses promote response repetition. Thus, although both studies show that experience influences subsequent behavior in teleosts, they reveal distinct forms of history dependence: automatic motor-response in zebrafish versus decision-making in archerfish.

Despite the insights gained into teleost decision-making, our study has certain limitations. The restricted nature of our behavioral paradigms may not reflect the full complexity of naturalistic choice, and our framework has yet to be integrated with neural recordings. Nevertheless, by illuminating these core choice determinants, we establish a vital behavioral baseline to guide future physiological investigations.

Together, these results demonstrate that key determinants of mammalian decision-making are shared by teleost fish, pointing to ancient, conserved principles of vertebrate choice that transcend fundamental differences in brain architecture and ecological niche. Concurrently, the persistence of archerfish choices during reward omission points to key divergences in how different taxa update choices under changing contingencies.

## MATERIALS AND METHODS

### Animals

A total of 18 archerfish (*Toxotes chatareus*) participated in this study, of which 14 completed the data collection stage and 4 died of natural causes. Adult fish measuring 6-13 cm in length and weighing 10-17 g were included. The archerfish were purchased from a local supplier and housed individually in 100 L aquaria containing brackish water. The water temperature was maintained at 25-29°C in a 12-hour light/dark cycle. Due to the absence of external sexual dimorphism in Toxotes species, individual sex could not be determined visually. Consequently, male and female subjects were pooled and analyzed as a single experimental group. Data collection continued for each subject for as long as it cooperated, after which it was returned to the stock tank containing fish not participating in the experiment. The number of trials collected for each fish ranged from 304 to 629, as summarized in Table 1.

**Table 1.** Total number of trials per fish.

| Fish | Number of trials |
| --- | --- |
| 1 | 509 |
| 2 | 588 |
| 3 | 599 |
| 4 | 578 |
| 5 | 589 |
| 6 | 366 |
| 7 | 536 |
| 8 | 585 |
| 9 | 325 |
| 10 | 492 |
| 11 | 510 |
| 12 | 400 |
| 13 | 629 |
| 14 | 304 |

All 14 subjects completed the study, i.e., there was no attrition. Fish care and all experimental procedures were approved by the Ben-Gurion University of the Negev Institutional Animal Care and Use Committee, in accordance with the regulations of the State of Israel.

### Training

After acclimation to the laboratory environment, naïve fish were trained to shoot at visual targets displayed on a computer monitor. The display consisted of a 21.5-inch BenQ VW2245-T monitor (BenQ Corp., Taiwan), positioned 35 ± 2 cm above the surface of the water. Training took place three to four times per week, and each session was composed of 30 trials.

The fish were first trained to shoot at a single target consisting of either a ladybug image or a red circle displayed on a white background. The target appeared at randomized locations on the screen. Immediately before target onset, a blinking black square was presented in the center of the screen for 3 s, to attract the fish’s attention. A trial was scored as successful if the fish hit the target on the screen within 30 s and was followed by delivery of a food pellet. If the fish did not respond within 30 s, the target disappeared and the next trial began after a 10 s break. Training continued until the fish reached a criterion of 80% successful hits within the 30 s response window.

Next, the fish were trained on a two-alternative, non-forced choice procedure. On this task, the fish could choose either one of two images and were rewarded regardless of which one they selected. Sessions of 30 trials were repeated over multiple days to familiarize the fish with two-choice trials and to ensure reinforcement learning for both images. Eleven fish were trained on non-abstract shapes, and three fish were trained on abstract shapes, as described in detail below. Fish were considered trained and ready for testing after demonstrating >90% engagement in two consecutive sessions. Training typically required 2-6 sessions for the non-abstract shapes and 4-7 sessions for the abstract shapes. Only subsequent experimental sessions were recorded and used for analysis.

### Stimuli

#### Shapes

The stimuli consisted of images of spiders and ants. These are shapes that archerfish encounter in their natural environment. Insect images were obtained from the Bug wood Images project at insectimages.org. The other stimuli were rendered from animated 3D models of one spider and one ant that were purchased under a standard license from the Sketchfab store at sketchfab.com. All images were preprocessed in MATLAB (MATLAB, RRID:SCR_001622). Backgrounds were removed, the stimuli were converted to grayscale using MATLAB rgb2gray, and each object was placed on a white background. Object area was defined as the number of pixels inside the object contour, and on each trial the size level was drawn from a discrete uniform set of five values. These levels corresponded to approximately 10,000 pixels, 50,000 pixels, 100,000 pixels, 200,000 pixels, and 300,000 pixels.

#### Abstract shapes

The stimuli consisted of a red circle and morphed triangle. In a pilot study, the morphing level of the triangle was adjusted so that the mean accuracy across five sessions was approximately 65%.

### Experiment 1: two-position task

Experiment 1 examined selections of a specific shape, either a spider or an ant. Five fish were tested. Each trial presented one spider image and one ant image in a two-alternative, non-forced choice procedure. A food pellet was only delivered if the fish selected the designated target stimulus. Individual fish completed this longitudinal study for as many sessions as their health permitted, with 30 trials per session, over 3-10 weeks, typically 3 sessions per week. This schedule was chosen to reduce the risk of overfeeding and to preserve motivation.

The stimulus set included 400 images grouped into 200 spider-ant pairs. On each trial, one pair was selected from this set. Across subjects, engagement ranged from ∼60% to 99% of all trials. Target identity was balanced across fish, with some fish trained to select spiders and others trained to select ants. Images varied in viewpoint, orientation, size, and contrast, and all presentation parameters were randomized. The target and the non-target appeared at two fixed locations throughout the study.

### Experiment 2: four-position task

Experiment 2 was identical to Experiment 1, except that target and non-target stimuli were randomly positioned anywhere on the screen. To maintain a sufficient distance between the two images, they were constrained from appearing in the same quadrant. This experimental design maintained adequate spatial separation while randomizing the rewarded location.

### Experiment 3: four-position task with abstract stimuli

Experiment 3 tested selection of an abstract shape. The stimulus pair was a red circle and a morphed red triangle. For each fish, one stimulus served as the target and the other as the non-target. The target and non-target locations were randomized across the screen, with the constraint that the two images could not appear in the same quadrant.

### Dynamic Generalized Linear Model

To characterize the trial-by-trial choices, we fitted a series of dynamic generalized linear models to each archerfish’s binary choice data [2]. In this framework, the probability of a correct response (choosing the target) on trial t was modelled as a logistic function of a linear combination of time-varying weights of the decision variables:

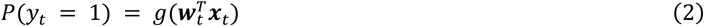

where *y_t_* = 1 denotes choosing the target on trial *t*, *g*(·) denotes the logistic function, ***x****_t_* is the vector of decision-making variables on trial *t*, and ***w****_t_* is the corresponding vector of weights. Vectors appear in bold and scalars are un-bolded. Each weight evolved across trials according to a Gaussian random walk prior:

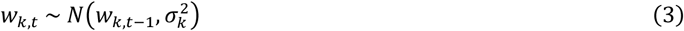

where *σ_k_*^2^ governed the volatility (rate of change) of the *k*-th weight across trials. This formulation allowed each weight to drift smoothly over time to capture gradual changes in the archerfish’s decision-making.

### Decision-Making Variable Definitions

The full form of the decision-making variables, ***x****_t_* in equation (2), is given by:

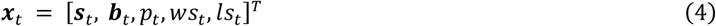

where ***s****_t_*, ***b****_t_*, denote the target position and spatial bias vectors on trial *t*, respectively, and *p_t_*, *ws_t_*, and *ls_t_* are the stimulus-history, win-stay, and lose-stay variables, respectively. The meaning and encoding of each variable are described below.

- Target position (***s****_t_*): A 2-element vector representing the position of the target on trial *t*. The *i*-th element (*i* = 1, 2), *s_t_^i^*, was a binary variable coded as +1 when the target appeared at that position, and 0 otherwise. The vector ***s****_t_* was generalized to a 4-element vector in the tasks with 4 possible positions.
- Spatial bias (***b****_t_*): A vector representing the possible positions for the bias on trial t. The *i*-th element (*i* = 1), *b_t_^i^*, was coded as +1 if the archerfish chose that position on trial t (regardless of whether it was the target), -1 if that position was available but not chosen on trial t, and 0 if that position was not presented. The vector ***b****_t_* was generalized to a 3-element vector on tasks with 4 possible positions. These weights captured the archerfish’s developing spatial aversions or spatial preferences.
- Stimulus-history (*p_t_*): Encoded as +1 when the archerfish chose the same shape on trial t as in the previous trial, t-1; e.g., a correct choice followed by a correct choice, or an incorrect choice followed by an incorrect choice, and -1 when the archerfish switched shapes between trial t-1 and trial t.
- Win-stay (*ws_t_*): Applicable only after correct trials; i.e., when the archerfish had successfully selected the target on the previous trial, t-1, it was coded as +1 and 0 otherwise. This variable captured the tendency to repeat a rewarded position.
- Lose-stay (*ls_t_*): The opposite of win-stay: this variable was applicable only after incorrect trials; i.e., when the archerfish had not selected the target on trial t-1, it was coded as +1, and 0 otherwise. This variable captured the tendency to remain at a location despite receiving no reward.

### Model variants

We evaluated several model variants, each incorporating a different subset of the decision-making variables in equation (4), to determine which model best accounted for the archerfish’s choices. As summarized in Table 2, we compared seven models for the two-position task (Experiment 1) and five models for the four-position task (Experiment 2 and Experiment 3).

**Table 2.**
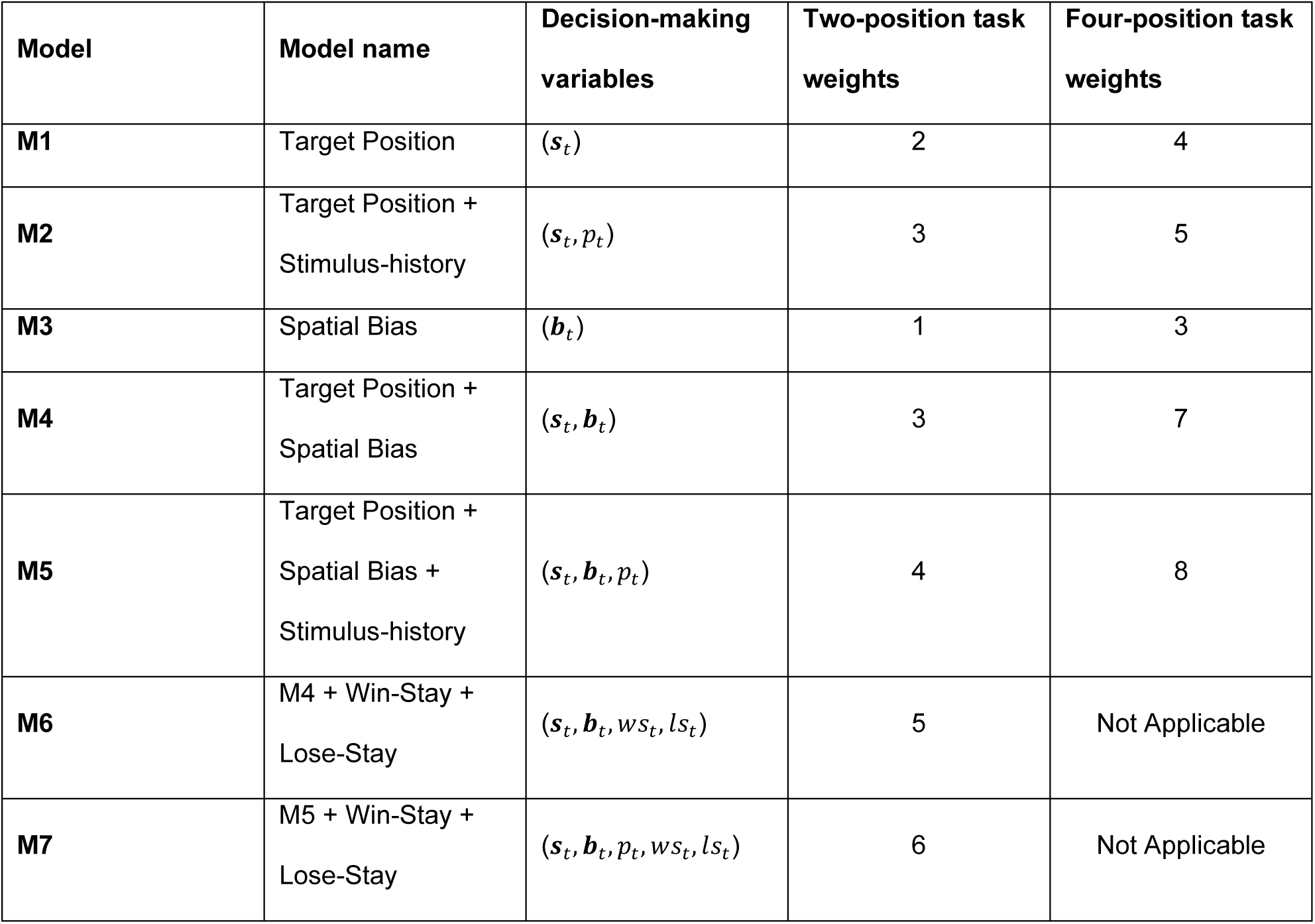
Summary of model parameters for the two-position and four-position task variants. The two rightmost columns indicate the number of weights used to model decision-making in each task.

| Model | Model name | Decision-making variables | Two-position task weights | Four-position task weights |
| --- | --- | --- | --- | --- |
| **M1** | Target Position | $(s_t)$ | 2 | 4 |
| **M2** | Target Position + Stimulus-history | $(s_t, p_t)$ | 3 | 5 |
| **M3** | Spatial Bias | $(b_t)$ | 1 | 3 |
| **M4** | Target Position + Spatial Bias | $(s_t, b_t)$ | 3 | 7 |
| **M5** | Target Position + Spatial Bias + Stimulus-history | $(s_t, b_t, p_t)$ | 4 | 8 |
| **M6** | M4 + Win-Stay + Lose-Stay | $(s_t, b_t, ws_t, ls_t)$ | 5 | Not Applicable |
| **M7** | M5 + Win-Stay + Lose-Stay | $(s_t, b_t, p_t, ws_t, ls_t)$ | 6 | Not Applicable |

### Inference algorithm

Model parameters were estimated using a decoupled Laplace empirical Bayes procedure [2]. The algorithm alternated between two steps. In the outer loop, the maximum a-posteriori (MAP) estimate of the weight trajectories was obtained via Newton’s method. Because multiple regressors could be nonzero on a single trial, the full negative Hessian of the log-posterior coupled all weights on each trial. Accordingly, we used a joint Newton step over all weights simultaneously by exploiting the block-tridiagonal structure of the Hessian (with *K x K* blocks) to obtain exact solutions via the block Thomas algorithm in time *O*(*Nk*³).

In the inner loop, the innovation variances *σ^2^_k_* of the random-walk prior (equation (3)) were optimized via the decoupled Laplace approximation. The Gaussian likelihood approximation was computed once from the current MAP estimate and held fixed, whereas the per-weight posterior precision matrices were updated using the closed-form sigma update: 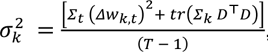, where Δ*w_k,t_* = *w_k,t_* – *w_k,t_* _–1_ denotes the first differences of the MAP weights, *Σ_k_* is the posterior covariance of the *k-th* weight trajectory, and *D* is the first-difference operator and *T* is the number of trials. The trace term accounted for posterior uncertainty in the weight trajectory and was computed via the tridiagonal inverse algorithm. Inner iterations continued until the relative change in all *σ_k_* values fell below 10^−4^.

The outer loop iterated up to 20 times, each time recomputing the MAP estimate and re-linearizing the likelihood before updating ***σ***. Convergence was assessed by monitoring the relative change in ***σ*** across outer iterations (tolerance: 10^−4^) and the maximum absolute change in weights during Newton iterations (tolerance: 10^−6^). The innovation variances were bounded between 10^−4^ and 100 to ensure numerical stability. The iteration yielding the highest marginal log-evidence (Laplace approximation) was retained as the final solution. Posterior credible intervals (±1.96 standard deviation) for each weight trajectory were obtained from the diagonal of the posterior covariance matrix and were computed via inversion of the tridiagonal posterior precision matrix at the converged MAP estimate.

### Model comparison and evaluation

The models were compared using the Bayesian Information Criterion, *BIC* = −2 log(*L*) + *K* log(*T*), where *L* is the maximum likelihood, *K* is the number of time-varying weight trajectories and *T* is the number of trials. The model with the lowest *BIC* was selected as the best-fitting model. The area under the receiver operating characteristic curve, *AUC*, was computed to assess each model’s discriminative ability to predict the archerfish’s binary choices. McFadden’s pseudo-*R*^2^ was calculated as 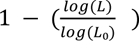 , where *L*₀ was the likelihood of a null model predicting the marginal choice probability on every trial. We also ran 10-fold cross-validation, in which the model was fit on a random 90% of the trials and tested on the remaining 10% at each iteration. Trial-level predictions were obtained by passing the linear predictor through the logistic function, *g* (equation (2)), and overall accuracy was computed as the proportion of trials on which the predicted choice matched the observed choice. Running accuracy was computed using a centered sliding window of ±15 trials (31 trials total), and smoothed model-fit overlays used a 20-trial moving average for visualization.

### Probe versus reinforced comparison

#### Design

For each fish we compared its performance on the probe trials with its performance on the reinforced trials. Sessions were run in repeating cycles of three: each cycle had one reinforcement session (R) and two probe sessions (P), and the reinforcement session shifted one step later on each cycle, which produced order R-P-P, then P-R-P, then P-P-R, before repeating. This rotation placed the reinforcement sessions at the start, middle, and end of the cycle equally often, to minimize the impact of a session’s position on results. In the probe sessions, the probe block was triggered by a correct response (this correct trial served as the block’s catch trial) and was positioned between trials 10 and 25.

#### Measures

Probe trials were defined as the responses within each “during-probe” block. By design, the first (catch) trial that was correct and with engagement was dropped. For the matched baseline, we used reinforced trials at similar within-session positions. To keep this baseline comparable to the probe blocks, reinforced trials were drawn solely from sessions in which the fish performed at criterion (at least 50% of trials completed). Probe sessions, which were only ever run when the fish was at criterion, were all kept. For each fish we then computed accuracy and engagement (mean ± SD across subjects), separately for the probe and reinforced trials and separately for the early and late phases of learning. The boundary between the early learning phase and late learning phase was the trial on which each fish attained the learning criterion, which was defined as the first trial where its smoothed accuracy (a centered 31-trial moving average) stayed above 65% correct for 30 consecutive trials. Fish 6 was only ever tested under reinforcement and was excluded.

#### Statistical analysis

We tested for similarity by computing each fish’s probe and reinforced accuracy and engagement and comparing the two conditions across fish with two one-sided tests (TOST), using an a-priori equivalence margin of ±0.15, the smallest difference in accuracy or engagement we considered meaningful for this task. Sample size (n=13) was sufficient, verified by power analysis with a level of significance of 0.05 and a power of 0.9.

All analyses were implemented in MATLAB R2024a, using custom code. The code for the analyses presented in this paper is openly accessible at https://doi.org/10.5281/zenodo.20679691.

## ACKNOWLEDGMENTS

This work was supported by the Israel Science Foundation (ISF) under Grant No. 824/21 (RS) and Grant No. 624/22 (MS).

## AUTHOR CONTRIBUTIONS

O.H. and R.S. conceptualized the study. O.H. and K.M. designed the experiments. O.H., K.M. and N.D.S. performed data collection. S.V. prepared the visual stimuli. O.H. and K.M. analyzed the data. O.H. and R.S. wrote the main manuscript text. O.H. and K.M. prepared figures 1-4. R.S. and M.S. jointly supervised the project. All authors reviewed and approved the final version of the manuscript.

## DATA AVAILABILITY

The datasets generated during and/or analyzed during the current study are available [60] in the Zenodo repository, https://doi.org/10.5281/zenodo.20679691

## ADDITIONAL INFORMATION

The authors declare no competing interests.

## Notes

https://zenodo.org/records/20679691

### Competing Interest Statement

The authors have declared no competing interest.

### Summary of Updates

This version of the manuscript has been revised to update the title, abstract, introduction, and discussion.

## REFERENCES

1. Busse, L. et al. The detection of visual contrast in the behaving mouse. J. Neurosci. 31, 11351–11361 (2011).

2. Roy, N. A., Bak, J. H., Akrami, A., Brody, C. D. & Pillow, J. W. Extracting the dynamics of behavior in sensory decision-making experiments. Neuron 109, 597–610 (2021).

3. Carandini, M. & Churchland, A. K. Probing perceptual decisions in rodents. Nat. Neurosci. 16, 824– 831 (2013).

4. Dayan, P. & Daw, N. D. Decision theory, reinforcement learning, and the brain. Cogn. Affect. Behav. Neurosci. 8, 429–453 (2008).

5. Urai, A. E., De Gee, J. W., Tsetsos, K. & Donner, T. H. Choice history biases subsequent evidence accumulation. eLife 8, e46331 (2019).

6. Kiani, R., Cueva, C. J., Reppas, J. B. & Newsome, W. T. Dynamics of neural population responses in prefrontal cortex indicate changes of mind on single trials. Curr. Biol. 24, 1542–1547 (2014).

7. Hwang, E. J., Dahlen, J. E., Mukundan, M. & Komiyama, T. History-based action selection bias in posterior parietal cortex. Nat. Commun. 8, 1242 (2017).

8. Akrami, A., Kopec, C. D., Diamond, M. E. & Brody, C. D. Posterior parietal cortex represents sensory history and mediates its effects on behaviour. Nature 554, 368–372 (2018).

9. Hanks, T. D. & Summerfield, C. Perceptual decision making in rodents, monkeys, and humans. Neuron 93, 15–31 (2017).

10. Venditto, S. J. C., Miller, K. J., Brody, C. D. & Daw, N. D. Dynamic reinforcement learning reveals time-dependent shifts in strategy during reward learning. eLife 13, RP97612 (2024).

11. Prétôt, L., Bshary, R. & Brosnan, S. F. Comparing species decisions in a dichotomous choice task: adjusting task parameters improves performance in monkeys. Anim. Cogn. 19, 819–834 (2016).

12. Funahashi, S. Physical features of visual images affect macaque monkey’s preference for these images. Front. Behav. Neurosci. 10, 212 (2016).

13. Bari, B. A. et al. Reinforcement learning modeling reveals a reward-history-dependent strategy underlying reversal learning in squirrel monkeys. Behav. Neurosci. 136, 46 (2022).

14. Ferrari-Toniolo, S., Bujold, P. M. & Schultz, W. Probability distortion depends on choice sequence in rhesus monkeys. J. Neurosci. 39, 2915–2929 (2019).

15. Lyamzin, D. R., Aoki, R., Abdolrahmani, M. & Benucci, A. Probabilistic discrimination of relative stimulus features in mice. Proc. Natl. Acad. Sci. USA 118, e2103952118 (2021).

16. Treviño, M. Stimulus similarity determines the prevalence of behavioral laterality in a visual discrimination task for mice. Sci. Rep. 4, 7569 (2014).

17. Treviño, M. et al. Controlled variations in stimulus similarity during learning determine visual discrimination capacity in freely moving mice. Sci. Rep. 3, 1048 (2013).

18. Lak, A. et al. Reinforcement biases subsequent perceptual decisions when confidence is low, a widespread behavioral phenomenon. eLife 9, e49834 (2020).

19. Wittmann, M. K. et al. Global reward state affects learning and activity in raphe nucleus and anterior insula in monkeys. Nat. Commun. 11, 3771 (2020).

20. Abrahamyan, A., Silva, L. L., Dakin, S. C., Carandini, M. & Gardner, J. L. Adaptable history biases in human perceptual decisions. Proc. Natl. Acad. Sci. USA 113, E3581–E3590 (2016).

21. Braun, A., Urai, A. E. & Donner, T. H. Adaptive history biases result from confidence-weighted accumulation of past choices. J. Neurosci. 38, 2418–2429 (2018).

22. Hermoso-Mendizabal, A. et al. Response outcomes gate the impact of expectations on perceptual decisions. Nat. Commun. 11, 1057 (2020).

23. Lak, A. et al. Dopaminergic and prefrontal basis of learning from sensory confidence and reward value. Neuron 105, 700–711 (2020).

24. Zhu, Z. & Kuchibhotla, K. V. Performance errors during rodent learning reflect a dynamic choice strategy. Curr. Biol. 34, 2107–2117 (2024).

25. López-Yépez, J. S., Martin, J., Hulme, O. & Kvitsiani, D. Choice history effects in mice and humans improve reward harvesting efficiency. PLoS Comput. Biol. 17, e1009452 (2021).

26. Butler, A. B. & Hodos, W. Comparative Vertebrate Neuroanatomy: Evolution and Adaptation (John Wiley & Sons, 2005).

27. Bräuer, J., Hanus, D., Pika, S., Gray, R. & Uomini, N. Old and new approaches to animal cognition: there is not “one cognition”. J. Intell. 8, 28 (2020).

28. Prétôt, L. et al. ManyFishes: a big team science collaboration on fish comparative cognition. Anim. Cogn. 29, 12 (2025).

29. O’Connell, L. A. & Hofmann, H. A. The vertebrate mesolimbic reward system and social behavior network: a comparative synthesis. J. Comp. Neurol. 519, 3599–3639 (2011).

30. Krause, M., Schulze, W. & Schuster, S. Learning and cognition in a decision made at reflex speed. eLife 13, e99634 (2024).

31. Ben-Simon, A., Ben-Shahar, O. & Segev, R. Measuring and tracking eye movements of a behaving archer fish by real-time stereo vision. J. Neurosci. Methods 184, 235–243 (2009).

32. Ben-Tov, M., Donchin, O., Ben-Shahar, O. & Segev, R. Pop-out in visual search of moving targets in the archer fish. Nat. Commun. 6, 6476 (2015).

33. Gabay, S. et al. Inhibition of return in the archer fish. Nat. Commun. 4, 1657 (2013).

34. Volotsky, S., Ben-Shahar, O., Donchin, O. & Segev, R. Recognition of natural objects in the archerfish. J. Exp. Biol. 225, jeb243237 (2022).

35. Volotsky, S., Donchin, O. & Segev, R. The archerfish uses motor adaptation in shooting to correct for changing physical conditions. eLife 12, RP92909 (2024).

36. Lau, B. & Glimcher, P. W. Dynamic response-by-response models of matching behavior in rhesus monkeys. J. Exp. Anal. Behav. 84, 555–579 (2005).

37. Mochol, G., Kiani, R. & Moreno-Bote, R. Prefrontal cortex represents heuristics that shape choice bias and its integration into future behavior. Curr. Biol. 31, 1234–1244 (2021).

38. Maoz, U. et al. Predeliberation activity in prefrontal cortex and striatum and the prediction of subsequent value judgment. Front. Neurosci. 7, 225 (2013).

39. Kim, Y. H., Park, J., Kim, Y. G., Lee, Y. & Kim, H. F. Value and spatial preferences guide habitual reaching and manual object selection in primates. Preprint at *bioRxiv* 10.64898/2026.06.20.733501 (2026).

40. Jahn, C. I. et al. Neural responses in macaque prefrontal cortex are linked to strategic exploration. PLoS Biol. 21, e3001985 (2023).

41. Goltstein, P. M., Reinert, S., Glas, A., Bonhoeffer, T. & Hübener, M. Food and water restriction lead to differential learning behaviors in a head-fixed two-choice visual discrimination task for mice. PLoS ONE 13, e0204066 (2018).

42. Dickson, P. E. & Mittleman, G. Visual discrimination, serial reversal, and extinction learning in the mdx mouse. Front. Behav. Neurosci. 13, 200 (2019).

43. Hachen, I., Reinartz, S., Brasselet, R., Stroligo, A. & Diamond, M. E. Dynamics of history-dependent perceptual judgment. Nat. Commun. 12, 6036 (2021).

44. Hughes, R. N. Lack of spontaneous alternation in favor of perseveration in domestic fowls and pigeons. Behav. Process. 20, 85–92 (1989).

45. Stagner, J. P., Michler, D. M., Rayburn-Reeves, R. M., Laude, J. R. & Zentall, T. R. Midsession reversal learning: why do pigeons anticipate and perseverate? Learn. Behav. 41, 54–60 (2013).

46. Karoubi, N., Segev, R. & Wullimann, M. F. The brain of the archerfish *Toxotes chatareus*: a Nissl-based neuroanatomical atlas and catecholaminergic/cholinergic systems. Front. Neuroanat. 10, 106 (2016).

47. Kuchibhotla, K. V. et al. Dissociating task acquisition from expression during learning reveals latent knowledge. Nat. Commun. 10, 2151 (2019).

48. Waiblinger, C., McDonnell, M. E., Reedy, A. R., Borden, P. Y. & Stanley, G. B. Emerging experience-dependent dynamics in primary somatosensory cortex reflect behavioral adaptation. Nat. Commun. 13, 534 (2022).

49. Li, J., Tang, L., Wei, X., Chen, Y. & Xu, H. Flexible goal learning involves coordinated population activity in dCA1 and medial orbitofrontal cortex. PLoS Biol. 24, e3003824 (2026).

50. Pisupati, S., Chartarifsky-Lynn, L., Khanal, A. & Churchland, A. K. Lapses in perceptual decisions reflect exploration. eLife 10, e55490 (2021).

51. Ashwood, Z. C. et al. Mice alternate between discrete strategies during perceptual decision-making. Nat. Neurosci. 25, 201–212 (2022).

52. Le, N. M. et al. Mixtures of strategies underlie rodent behavior during reversal learning. PLoS Comput. Biol. 19, e1011430 (2023).

53. Mackintosh, N. J. & Cauty, A. Spatial reversal learning in rats, pigeons, and goldfish. Psychon. Sci. 22, 281–282 (1971).

54. Parker, M. O. et al. Discrimination reversal and attentional sets in zebrafish (*Danio rerio*). Behav. Brain Res. 232, 264–268 (2012).

55. Urai, A. E. & Donner, T. H. Persistent activity in human parietal cortex mediates perceptual choice repetition bias. Nat. Commun. 13, 6015 (2022).

56. Rink, E. & Wullimann, M. F. The teleostean (zebrafish) dopaminergic system ascending to the subpallium (striatum) is located in the basal diencephalon (posterior tuberculum). Brain Res. 889, 316– 330 (2001).

57. Tay, T. L., Ronneberger, O., Ryu, S., Nitschke, R. & Driever, W. Comprehensive catecholaminergic projectome analysis reveals single-neuron integration of zebrafish ascending and descending dopaminergic systems. Nat. Commun. 2, 171 (2011).

58. Amo, R. et al. The habenulo-raphe serotonergic circuit encodes an aversive expectation value essential for adaptive active avoidance of danger. Neuron 84, 1034–1048 (2014).

59. Mangalwedhekar, A., Hunt, S., Hachen, I. & Bahl, A. Opposing influences of sensory and response history in larval zebrafish. Preprint at *bioRxiv* 10.64898/2026.07.30.741669 (2026).

60. Hendler, O. Code and data for manuscript: Decision-making dynamics mask the true psychophysical capacity of archerfish. Zenodo 10.5281/zenodo.20679691 (2026).

