## Supplementary material for "Trial-History and Spatial Biases Shape Perceptual Decision-Making in Archerfish": S1 - S5

### Supplementary materials

#### Supplementary Figure 1 (S1). Hendler, Mondal et al.

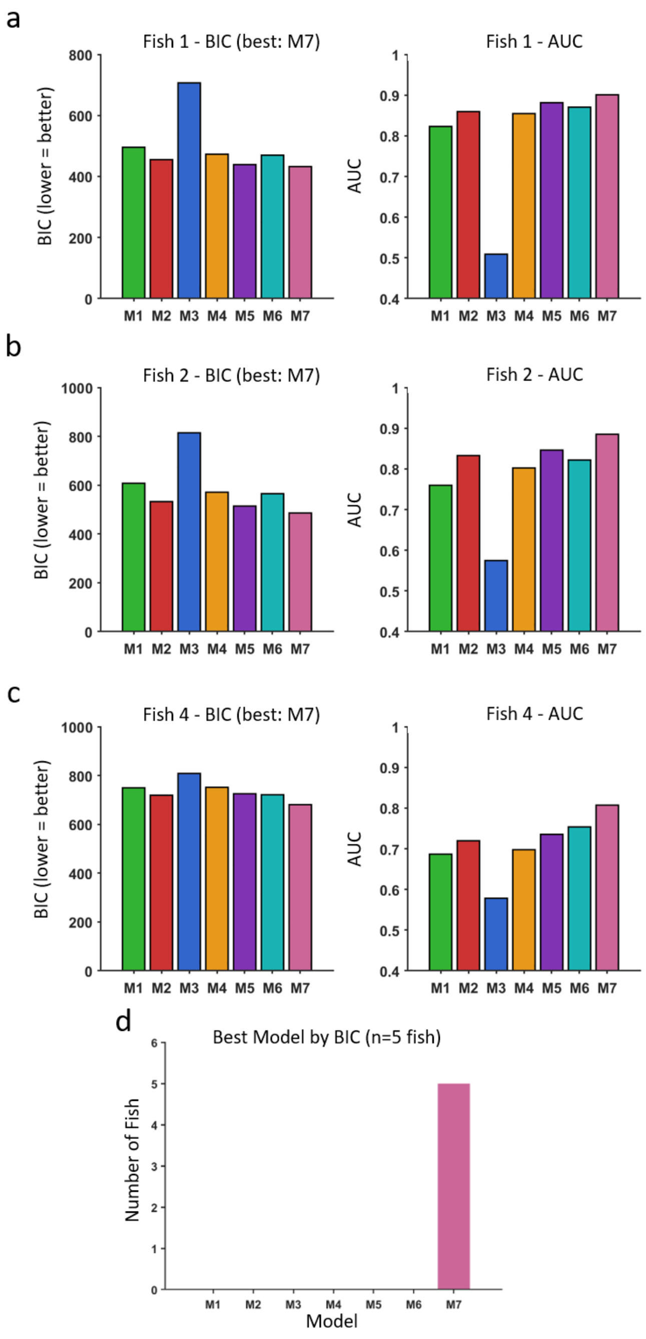


*Supplementary Material 1 (S1).* **BIC and AUC for candidate Models, two-position task**

**(a–c)** Bar plots showing the BIC (left) and AUC (right) for candidate models M1-M7, for Fish 1, Fish 2 and Fish 4, respectively. **(d)** Number of fish for which each model achieved the lowest BIC; M7 was the best-fitting model for all five fish in the two-position task.

### Supplementary Figure 2 (S2). Hendler, Mondal et al.

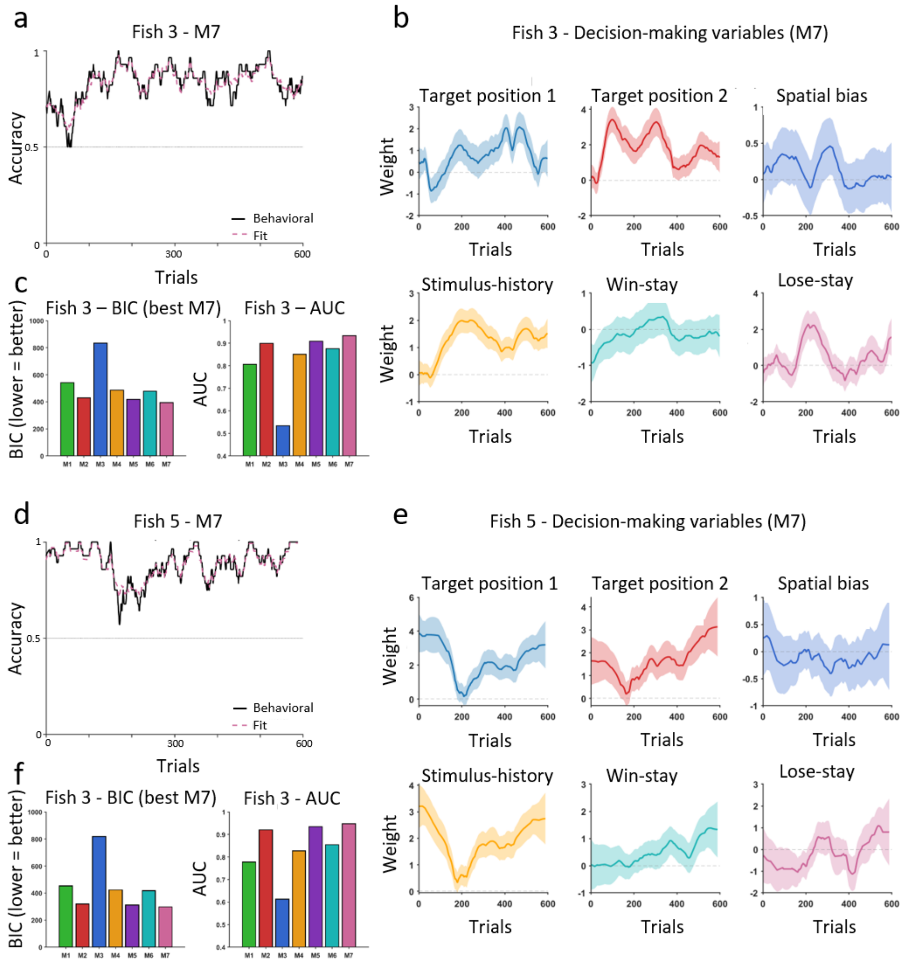


*Supplementary Material 2 (S2).* **Model fit and decision variables for Experiment 1, Fish 3 and Fish 5**

**(a)** Behavioral accuracy over trials for Fish 3. Accuracy as a function of trial number is depicted in black and the best-fitting model, M7, appears in dashed purple. The horizontal dashed grey line marks the chance value of 0.5. **(b)** Top row depicts the weights of the three spatial decision-making variables for Fish 3; i.e., target position 1, target position 2, and spatial bias. Solid lines show the posterior MAP estimates. Shaded bands represent ±1.96 posterior SD. Bottom row depicts the weights of the three history decision variables for the fish; i.e., stimulus-history, win-stay and lose-stay. **(c)** Comparison of all seven candidate models, M1–M7, indicating BIC and AUC for each. M7 had the lowest BIC and the highest AUC. **(d-f)** The same as panel A to C for Fish 5.

### Supplementary Figure 3 (S3). Hendler, Mondal et al.

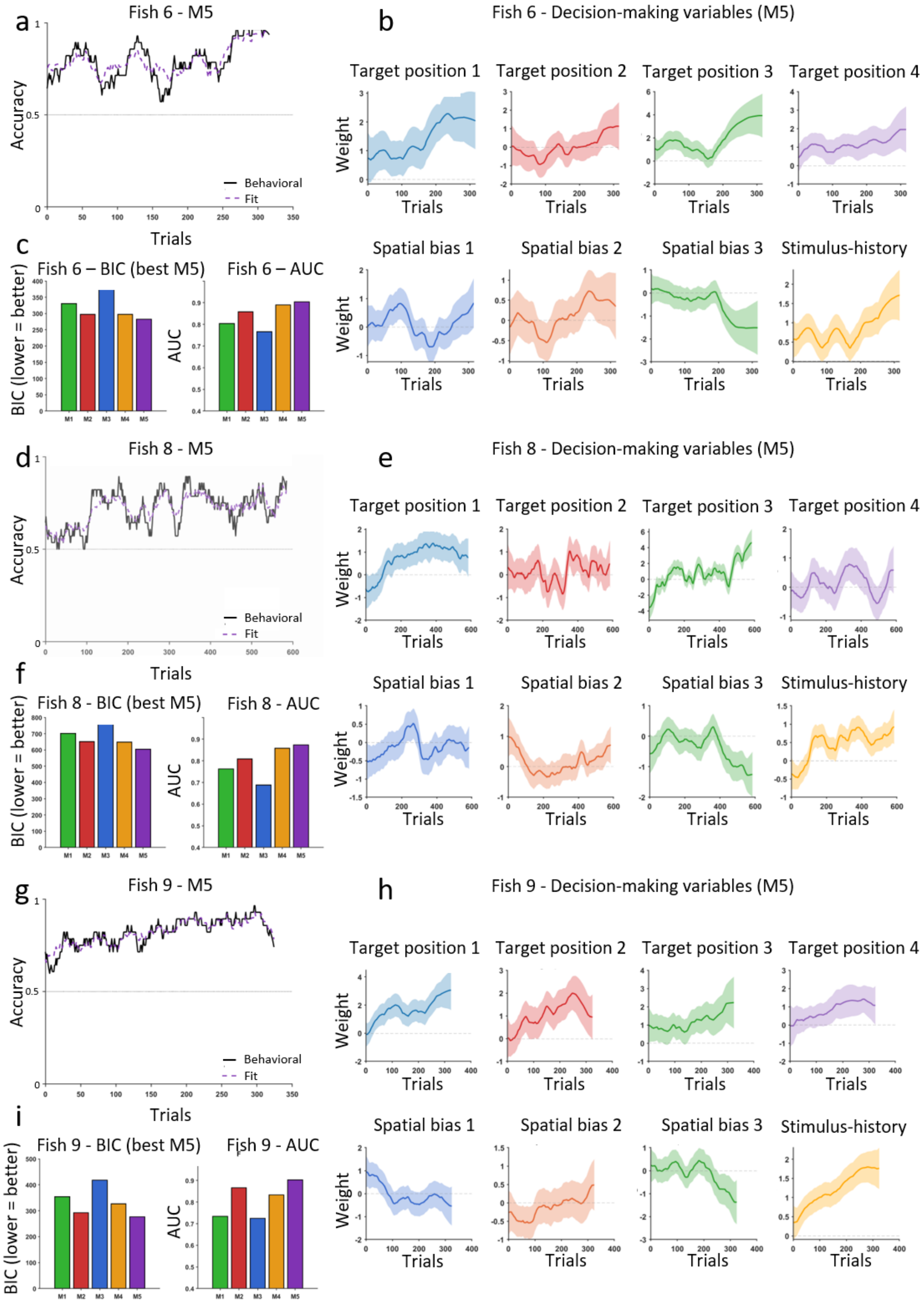


*Supplementary Material 3 (S3).* **Model fit and decision variables for Experiment 2, Fish 6, 8, 9**

**(a)** Behavioral accuracy as a function of trial number for Fish 6. Accuracy is shown in black and the best-fitting model M5 appears in dashed purple. Chance value is denoted by a gray dashed line. **(b)** Top row shows the weights for the four target position decision-making variables for Fish 6; i.e., target position 1, target position 2, target position 3 and target position 4. Solid lines show the posterior MAP estimates and shaded bands represent the ±1.96 posterior SD. Bottom row shows the weights for the three spatial bias variables for the fish: spatial bias 1, spatial bias 2, and spatial bias 3 and as well as stimulus-history. **(c)** Shows the comparison across all candidate models M1-M5, BIC for the five candidate models M1-M5 followed by the AUC for each model. M5 had the lowest BIC and the highest AUC across all models. **(d-f)** The same as panel A to C for Fish 8. **(g-i)** The same as panel A to C for Fish 9.

Supplementary Figure 4 (S4). Hendler, Mondal et al.

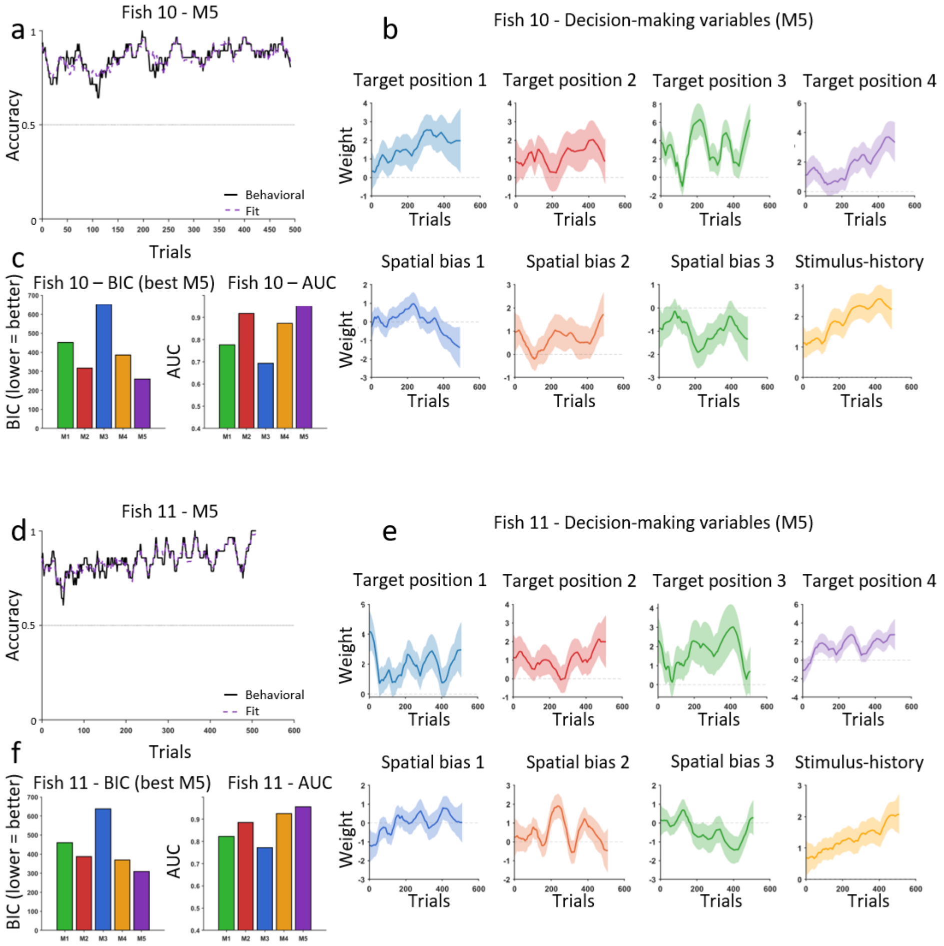


*Supplementary Material 4 (S4).* **Model fit and decision variables for Experiment 2, Fish 10 and Fish 11**

**(a)** Behavioral accuracy over trials for Fish 10. Black line represents accuracy as a function of trial number and dashed purple depicts the best-fitting model M5. The horizontal dashed grey line marks the chance value. **(b)** Top row displays the weights for four target position decision-making variables for Fish 10: target position 1, target position 2, target position 3 and target position 4. Solid lines show the posterior MAP estimates and shaded bands represent the ±1.96 posterior SD. Bottom row displays the weights for three spatial bias variables for the fish; i.e., spatial bias 1, spatial bias 2, and spatial bias 3 and the weight of stimulus-history. **(c)** Comparison of all candidate models, BIC for the five candidate models M1-M5 followed by the AUC for each model. M5 had the lowest BIC and the highest AUC of all models. **(d-f)** The same as panel A to C for Fish 11.

### Supplementary Figure 5 (S5). Hendler, Mondal et al.

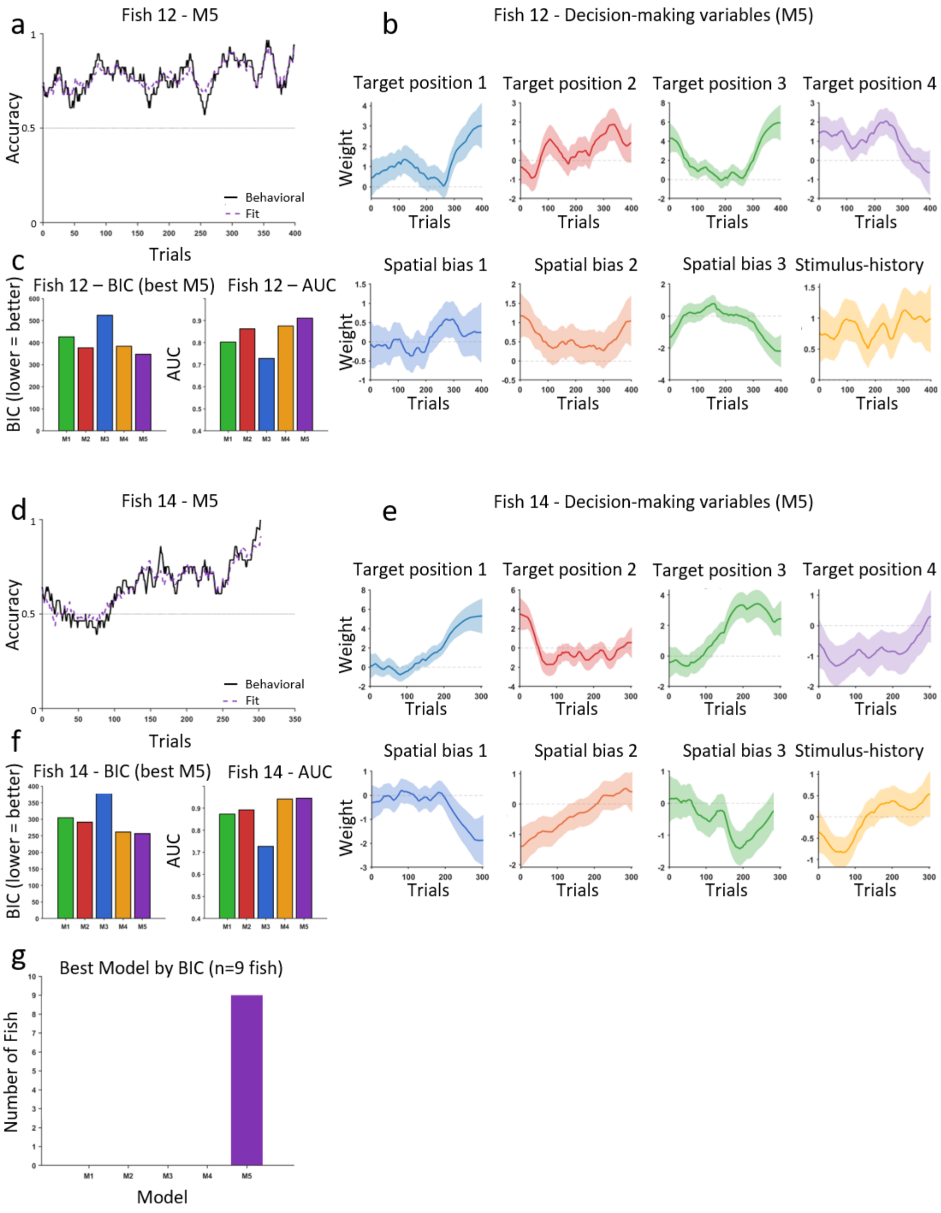


*Supplementary Material 5 (S5).* **Model fit and decision variables for Experiment 3, Fish 12 and Fish 14**

**(a)** Behavioral accuracy across trials for Fish 12. Accuracy is displayed in black and the best-fitting model M5 in dashed purple. The horizontal dashed grey line shows the chance value of 0.5. **(b)** Top row shows the weights for the four target position decision-making variables for Fish 6: target position 1, target position 2, target position 3 and target position 4. Solid lines show the posterior MAP estimates and the shaded bands represent the ±1.96 posterior SD. Bottom row depicts the weights of the three spatial bias variables for the fish: spatial bias 1, spatial bias 2, and spatial bias 3 and the weight of stimulus-history. **(c)** Comparison across all candidate models, the BIC for the five candidate models M1-M5 followed by the AUC for each model. M5 had the lowest BIC and highest AUC across all models. **(d-f)** The same as panel A to C for Fish 14. **(g)** The full model M5 which combined target position, spatial bias, stimulus-history, win-stay, and lose-stay was the best-fitting model for each fish.
